# BLAST-based validation of metagenomic sequence assignments

**DOI:** 10.1101/181636

**Authors:** Adam L. Bazinet, Brian D. Ondov, Daniel D. Sommer, Shashikala Ratnayake

## Abstract

When performing bioforensic casework, it is important to be able to reliably detect the presence of a particular organism in a metagenomic sample, even if the organism is only present in a trace amount. For this task, it is common to use a sequence classification program that determines the taxonomic affiliation of individual sequence reads by comparing them to reference database sequences. As metagenomic data sets often consist of millions or billions of reads that need to be compared to reference databases containing millions of sequences, such sequence classification programs typically use search heuristics and databases with reduced sequence diversity to speed up the analysis, which can lead to incorrect assignments. Thus, in a bioforensic setting where correct assignments are paramount, assignments of interest made by “first-pass” classifiers should be confirmed using the most precise methods and comprehensive databases available. In this study we present a blast-based method for validating the assignments made by less precise sequence classification programs, with optimal parameters for filtering of blast results determined via simulation of sequence reads from genomes of interest, and we apply the method to the detection of four pathogenic organisms. The software implementing the method is open source and freely available.

## Introduction

In metagenomic analysis, comparing the genomic sequence content of a sample to a reference database is fundamental to understanding which organisms present in the sample were sequenced. There exist many bioinformatics software programs that perform this classification task [7, 10, 22, 32]; some programs only estimate overall taxonomic composition and abundance in the sample [19,30], while other programs assign a taxonomic label to each metagenomic sequence [4, 14, 15, 17, 18, 28, 36]. In a bioforensic setting, one is often concerned with reliably detecting the presence of a particular organism in a metagenomic sample, which may only be present in a trace amount. For this task, one typically uses the latter class of programs just mentioned, which determine the taxonomic affiliation of each sequence using a reference database [8, 20, 21] and a taxonomy [5]. A canonical metagenomic sequence classification workflow is shown in Figure 1.

**Figure 1.**
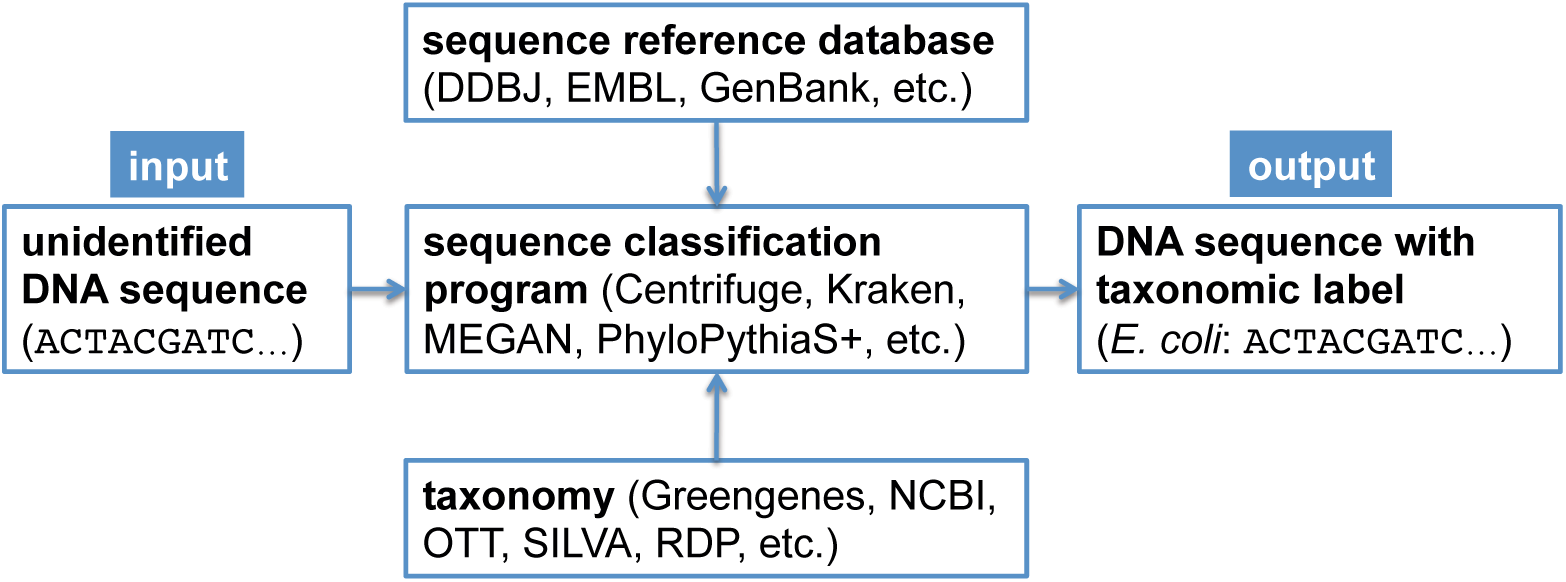
Canonical workflow for the classification of metagenomic sequences. A sequence classification program, which typically makes use of a reference database and a taxonomy, is used to assign taxonomic labels to unidentified dna sequences.

When classifying sequences, there is a general trade-off between sensitivity (the proportion of the total number of sequences assigned correctly) and precision (the proportion of assigned sequences assigned correctly), as well as between classification performance (combined sensitivity and precision) and computational resource requirements. Modern metagenomic sequence classification programs often use relatively fast heuristics and databases with limited sequence diversity to increase analysis speed, as metagenomic data sets often consist of millions or billions of sequences that need to be compared to millions of database sequences. Thus, while they are useful in performing a “first-pass” analysis, in a bioforensic setting it is important to validate the assignments of interest made by such programs using the most precise methods available [13]. One could choose to validate only the assignments made to the taxonomic clade of interest (e.g., *Bacillus anthracis)*, but depending on the compute capacity one has access to, one might choose to validate all assignments subsumed by a higher ranking taxon (e.g., the *Bacillus cereus* group or the *Bacillus* genus), which would enable the detection of possible false negative assignments as well as false positive assignments made by the first-pass classifier.

In this study, we present a method that uses blast [11], the ncbi non-redundant nucleotide database [12] (nt), and the ncbi taxonomy [12] to validate the assignments made by less precise sequence classification programs. blast is widely considered the “gold standard” for sequence comparison, although it is generally known to be orders of magnitude slower than the most commonly used first-pass classifiers (see Bazinet and Cummings, 2012 for a comparison of blast runtimes to those of other sequence classification programs). For simplicity, we refer to the taxonomic clade of interest in our analyses as the “target taxon”, and we assume all metagenomic sequences are paired-end reads generated by the Illumina HiSeq 2500 sequencer (no assembled sequences). The basic validation procedure involves comparing each read against the nt database using blastn, and then filtering and interpreting the blast results based on data collected from simulated read experiments aimed at optimising detection of the target taxon. The blast-based validation workflow is shown in Figure 2.

**Figure 2.**
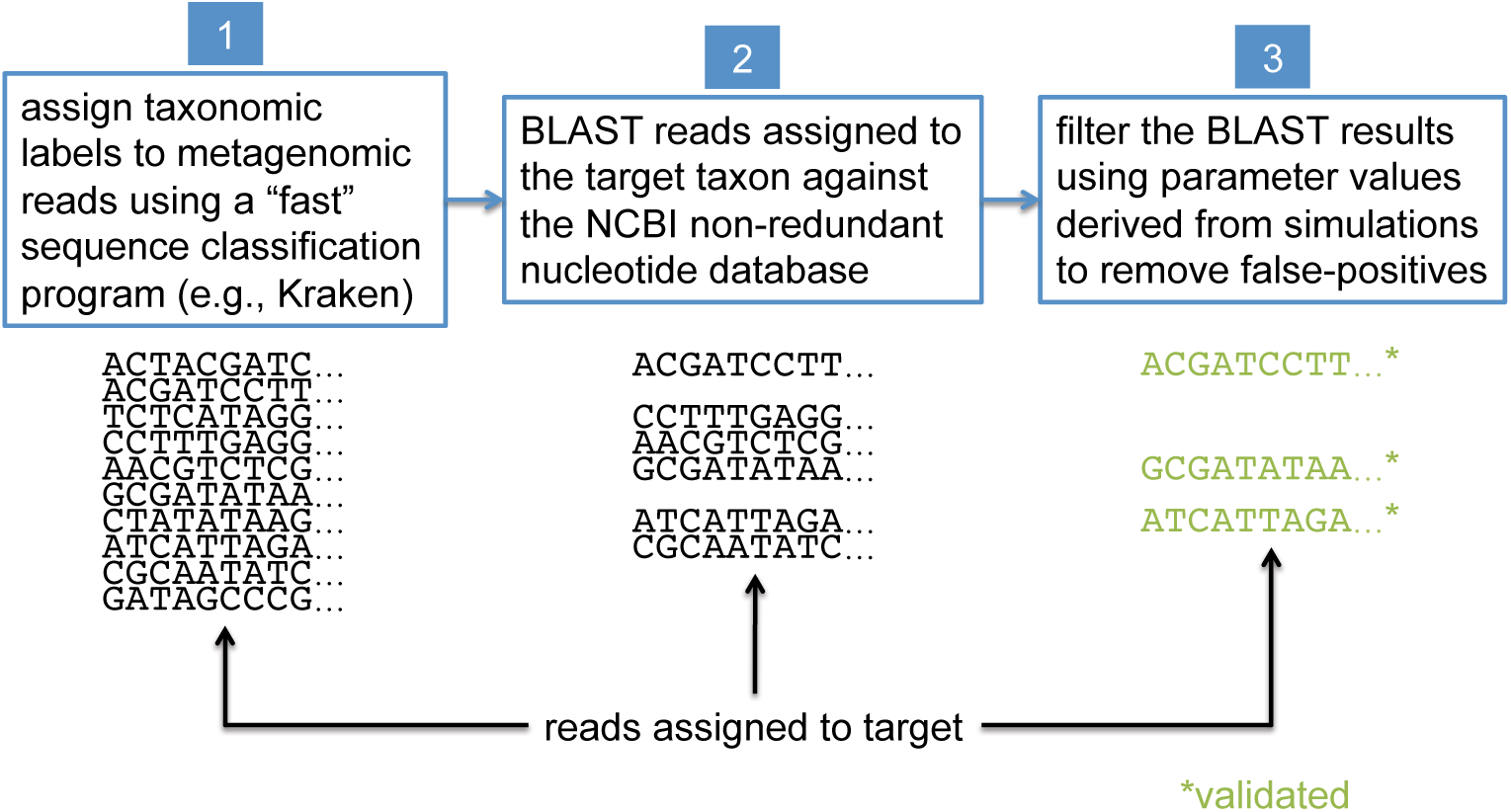
Workflow for blast-based validation of taxonomic assignments. Taxonomic labels are first assigned to metagenomic reads using a “first-pass” classification program. Reads assigned to a target taxon of interest are then compared against the ncbi nt database using blast. Final taxonomic assignments are obtained by filtering the blast results using parameter values that were previously determined to be optimal for the target taxon.

## Related work

### Platypus Conquistador

An existing software tool, “Platypus Conquistador” [13], also uses blast to validate the classification of particular sequences of interest. Platypus requires the user to split their reference sequences into two databases: a database containing only sequences of possible interest, and a database composed of potential “background” sequences. blast queries are performed against both databases, and hits may be filtered by various combinations of percent identity or alignment length values, which need to be provided by the user. After filtering, query sequences with hits to the “interest” database are checked to see if they also have hits to the “background” database; if so, the bit scores of the respective best hits are compared and are roughly categorised as “equal”, “interest > background”, etc. While this could be a helpful diagnostic tool, there is no guidance provided to the user as to what parameter values to use or what difference in bit scores between interest and background should be regarded as significant. Furthermore, this tool no longer appears to be actively developed.

### Genomic purity assessment

Whereas in this study we are concerned with the precision with which individual reads are classified so as to be confident in the detection of a target taxon in a metagenomic sample, a recently published study [27] addresses a different, but related problem, namely detecting contaminant organisms in ostensibly axenic (non-metagenomic) samples. Specifically, Olson et al. develop methods to determine the proportion of a contaminant required to be present in an otherwise pure material such that the contaminant can be detected with standard metagenomic sequence classification tools. As in our study, they simulate reads with art [16] software (in their case from both “material” and potential contaminant genomes) to set up conditions under which sequence classification performance can be assessed. PathoScope [15] is used instead of blast for read classification. In general, they find that their method is able to identify contaminants present in a proportion of at least 10^−3^ for most contaminant-material combinations tested.

### Outlier detection in BLAST hits

Shah et al. [33] have developed a method that detects outliers among blast hits in order to separate the hits most closely related to the query from hits that are phylogenetically more distant using a modified form of Bayesian Integral Log Odds (bild) scores [3] and a multiple alignment scoring function. In this way, they separate sequences with confident taxonomic assignments from sequences that should be analyzed further. The method was developed for and tested on 16S rrna data, and thus is currently not applicable to whole genome sequencing (wgs) data sets. As a general-purpose filter, however, it can be used with any organism containing 16S rrna data, whereas our methods are optimised for detection of specific taxa. It is also interesting to note that in the Shah et al. study, blast is used as a first-pass classifier and subsequent analysis is performed with tipp [23], whereas in the paradigm we present here, a much faster classifier than blast would be used for a first-pass (e.g., Kraken [36]), and then our blast-based method would be used for validation.

## Methods

### Evaluation of a “first-pass” taxonomic classifier

To demonstrate typical use of a first-pass taxonomic classification program, we used Kraken [36] (version 1.1). Kraken was run in paired-end mode with default parameters and used standard Kraken databases for bacteria, archaea, viruses, plasmids, and human sequences.

### Read simulation

For read simulation we used ART [16] (version 2016-06-05). To ensure thorough sampling, all experiments used simulated reads equivalent in total to 10× coverage of the source genome. For 150-bp reads, we used the built-in HS25 quality profile with an insert size of 200 ± 10 bp (mean ± standard deviation). For 250-bp reads, we used a custom Illumina HiSeq quality profile that we generated from recent runs of our HiSeq 2500, with an insert size of 868 ± 408 bp determined from recent library preparations. We supplied this information so that the simulated reads would have characteristics that closely matched what we would expect to obtain from a real HiSeq run in our laboratory, thus ensuring that the simulation results would be maximally useful to us. We recommend that others who emulate our procedures customise the attributes of their simulated reads to correspond to the real data they anticipate analysing.

### Similarity searches

We used blastn from the blast+ package [11] (version 2.2.25+) together with the ncbi non-redundant nucleotide database [12] (downloaded February 2017) for all classification experiments. Default parameters were used, except when excluding taxa from the reference database, in which case the -negative_gilist option was added. The blast computation was distributed over many cluster nodes to complete the analyses in a timely manner.

### BLAST result filters

The output from blast includes a number of statistics that can potentially be used to filter the results, including alignment length, alignment percent identity, E-value (the number of similar scoring alignments one can “expect” to see by chance in a database of the size being searched), and bit score (a database size-independent measure of alignment quality).

We developed two basic ways of filtering blast results. The first we term an “absolute” filter, which simply removes blast hits that do not meet a particular criterion. Various possible criteria include minimum alignment length, minimum alignment percent identity, or maximum E-value. Of these three filters, this study only uses the E-value filter (abbreviated *E*), as E-value is fundamentally a composite of alignment length and alignment similarity. (Our software supports the use of all three filters, however, either individually or in combination.) If the best blast hit matches the target taxon after application of the absolute filters, it is then possible to apply a “relative” filter by computing the difference in E-value or bit score between the best hit and the best hit to a *non-target* taxon (should the latter exist). As very small E-values are typically rounded to zero, our software uses relative bit scores in this context for maximum applicability; we call this quantity the “bit score difference”, abbreviated *b*. If *b* is greater than or equal to a threshold determined via read simulation experiments, then we have validated the assignment of the read to the target taxon. Examples of the application of the bit score difference filter are given in Figures 3 and 4.

**Figure 3.**
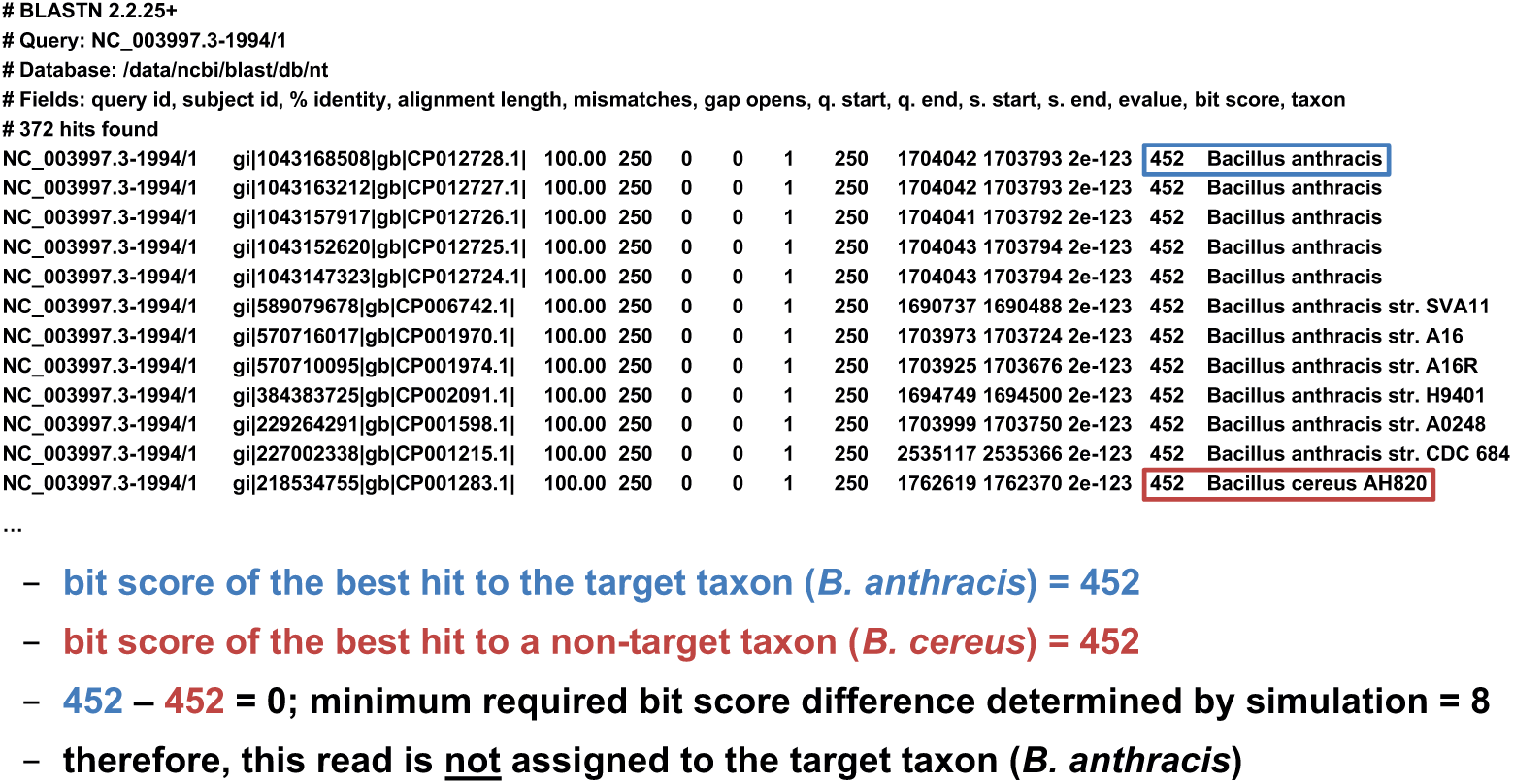
Demonstration of the “bit score difference” filter. In this first example, application of the bit score difference filter does not result in the assignment of the read to the target taxon.

**Figure 4.**
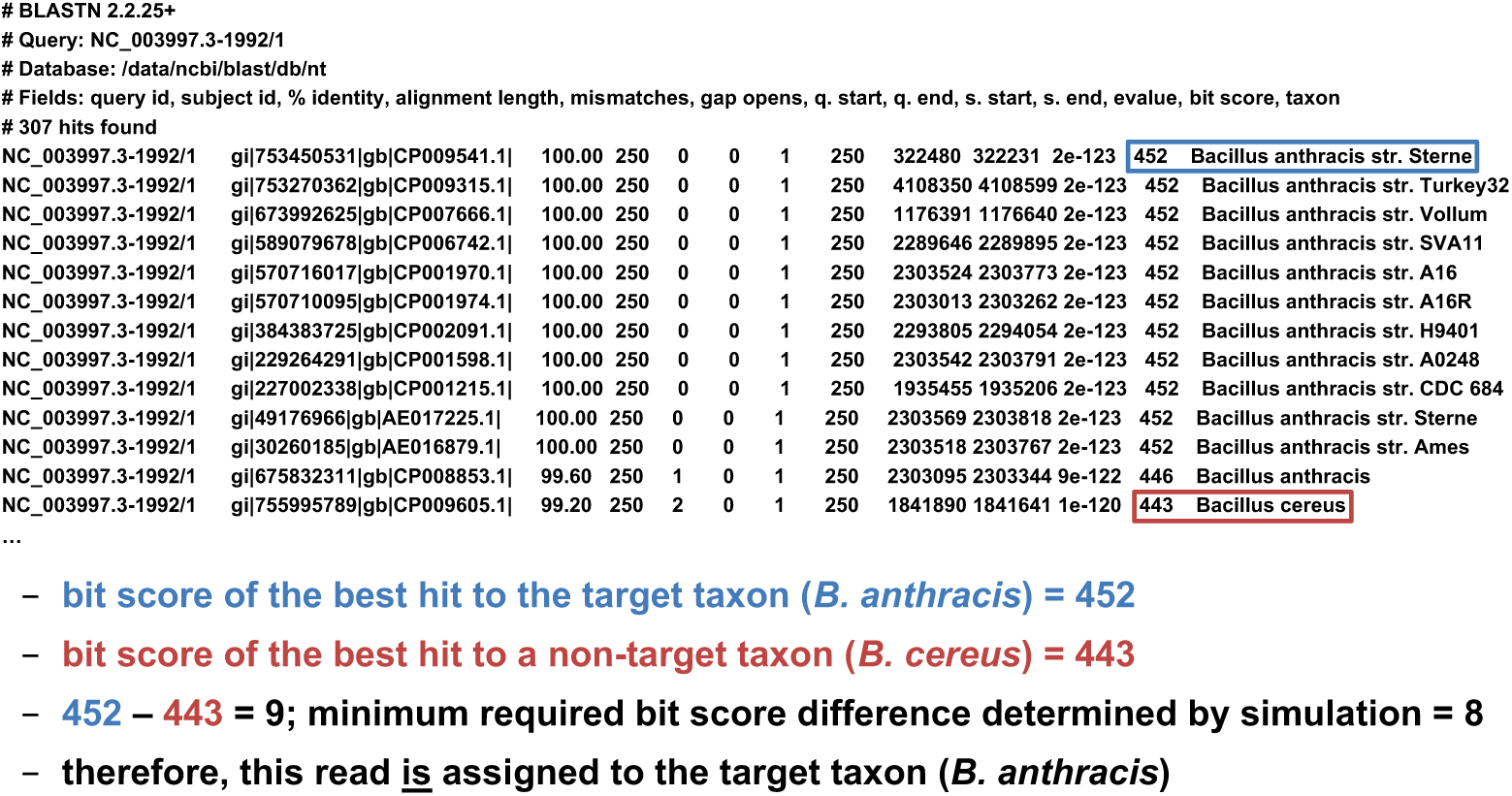
Demonstration of the “bit score difference” filter. In this second example, application of the bit score difference filter results in the assignment of the read to the target taxon.

### Evaluation of classification performance

The two main metrics used in this study to evaluate classification performance are sensitivity and precision.

To calculate sensitivity, one must determine the number of target taxon reads that were correctly assigned as a fraction of all the target taxon reads that were assigned. In this study, a true positive (*TP*) is a simulated read from the target taxon assigned correctly (either assigned directly to the target taxon or to a more specific taxon beneath the target), and a false negative (*FN*) is a simulated read from the target taxon assigned incorrectly (i.e., assigned to a taxon that is not part of the target taxon lineage). Note that the case of a non-specific but not incorrect read assignment (e.g., a *B. anthracis* read assigned to the *B. cereus group)* is neither considered a *TP* nor a *FN*; we term this an “inconclusive assignment” (*IA*). The count of true positives, false negatives, and inconclusive assignments can be easily determined by parsing the blast output associated with the target taxon. In all of our read simulation experiments, therefore, the calculation of sensitivity uses the formula (*TP*/(*TP* + *FN* + *IA*)).

To calculate precision, one must determine the number of non-target taxon reads incorrectly assigned to the target taxon, each of which is considered a false positive (*FP*). Naively, determining the count of false positives would require simulating reads and evaluating blast results for every non-target taxon in the database, but we currently regard this as computationally prohibitive. Instead, we offer two alternatives. The first, which we call “near neighbour”, computes *FP* using the genome in the database that is most globally similar to the target taxon as a proxy for all non-target database taxa. The intuition behind this approach is that a misclassified read (presumably due to sequencing error) is most likely to originate from a database genome that is very similar to the target taxon. Thus, with the near neighbour approach, the calculation of precision uses the formula (*TP*/(*TP* + *FP*)). The potential weakness of this approach is that there could be a region of local similarity to the target taxon in a database genome that is not the near neighbour. Thus, we offer a second approach that does not rely on selecting other genomes from the database, which we call the “false negatives” approach. This approach relies on the observation that if the sequencing error process is symmetric — i.e., the probability of an erroneous A to C substitution is the same as that of C to A, insertions are as probable as deletions, and so on — then the process that gives rise to false negatives can be treated as equivalent to the process that gives rise to false positives. While it is known that in practice this assumption of symmetry is violated [31], it may nonetheless suffice to use *FN* as a proxy for *FP* in this context. Thus, with the false negatives approach, it is only necessary to simulate reads from the target taxon, and the calculation of precision uses the formula (*TP*/(*TP* + *FN*)). Unfortunately, deciding which of the two heuristics is more effective would require comparison to a provably optimal procedure; in this study, we present results from simulated read experiments using both the near neighbour and false negatives approaches, and report the patterns we observe.

### BLAST result parsing, final taxonomic assignment, and calculation of statistics

blast result parsing and final taxonomic assignment of each read was performed with a custom Perl script capable of querying the ncbi taxonomy database [12]. If a target taxon is supplied as an argument to the script, assignments to the target taxon lineage that are more specific than the target taxon are simply reassigned to the target taxon. blast hits that do not meet the criteria specified by the absolute filters (minimum alignment length, minimum alignment percent identity, or maximum E-value) are removed, as are hits to the “other sequences” clade (ncbi taxon ID 28384), which are presumed to be erroneous. To make the final taxonomic assignment for each read, the lowest common ancestor (lca) algorithm [17] is applied to the remaining hits that have a difference in bit score from the best hit less than a specified amount. If multiple parameter values are supplied for one or more filters, the script parses the blast results once for each possible combination of parameter values and writes the results to separate “lca files”, thus enabling the user to efficiently perform parameter sweeps. The ultimate output from the script is one or more lca files, each containing the final taxonomic assignment of each read for a particular combination of filter parameter values. Counts of true positives, false negatives, inconclusive assignments, and false positives (from which sensitivity and precision were calculated) were obtained using a separate Perl script that parses the lca files produced by the blast result parser.

### Determination of optimal BLAST filter parameter values

When deciding how the absolute and relative BLAST filters should be parameterised, an optimality criterion is needed. In the execution of bioforensic casework, it is important that any assignments made are correct. Thus, we first chose filter parameter values that maximized precision (i.e., minimized incorrect read assignments). In the event that multiple combinations of parameter values yielded exactly the same maximum precision value, we chose from among these the combination that maximized sensitivity (i.e., maximized detection of the target taxon). In the event that multiple combinations of parameter values yielded exactly the same maximum precision *and* sensitivity values, we reported the strictest combination.

In this study, we present examples aimed at detecting a variety of pathogenic target taxa including *Bacillus anthracis, Clostridium botulinum*, pathogenic *Escherichia coli*, and *Yersinia pestis*. Due to inherent variation in the degree of interrelatedness among genomes from different taxonomic clades, optimal filter parameter values need to be set differently for each target taxon. To determine optimal filter settings, we must know the true origin of our test sequences; thus, we simulate reads from each target taxon genome, blast them against nt, and evaluate classification performance under different combinations of filter parameter settings. In each read simulation experiment, 81 different combinations of filter parameter values were tested — i.e., all combinations of maximum E-value (*E*) = {10^0^, 10^−1^, 10^−2^, 10^−4^, 10^−8^, 10^−16^, 10^−32^, 10^−64^, 10^−128^} and bit score difference (*b*) = {0, 1, 2, 4, 8, 16, 32, 64, 128}. The parameter optimisation workflow is shown in Figure 5.

**Figure 5.**
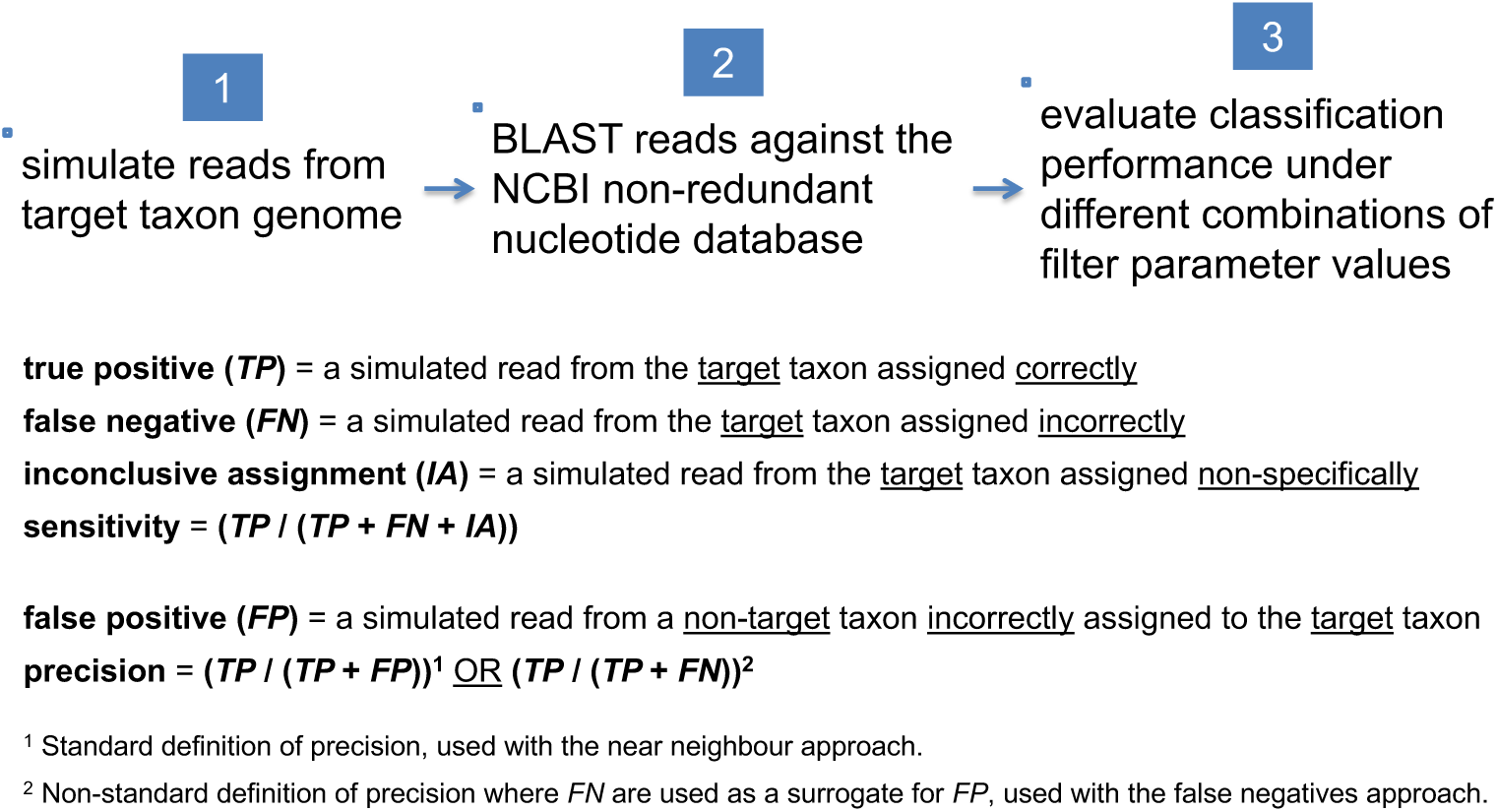
Workflow for determining optimal blast filter parameter values. Simulated reads from the target taxon genome are compared against the ncbi nt database using blast, and classification performance is evaluated under different combinations of parameter values used to filter blast results.

### Selection of near neighbour and alternate representative genomes

For each target taxon, we used the “Genome neighbor report” feature of the ncbi Genome database [12] to select the most closely related complete genome of a different species or strain, as appropriate, to be used as the “near neighbour”. For species-level target taxa, we used the Genome neighbor report to select the complete genome of the same species that was most distantly related to the original representative genome, which we call the “alternate representative genome” (Table 1).

**Table 1.**
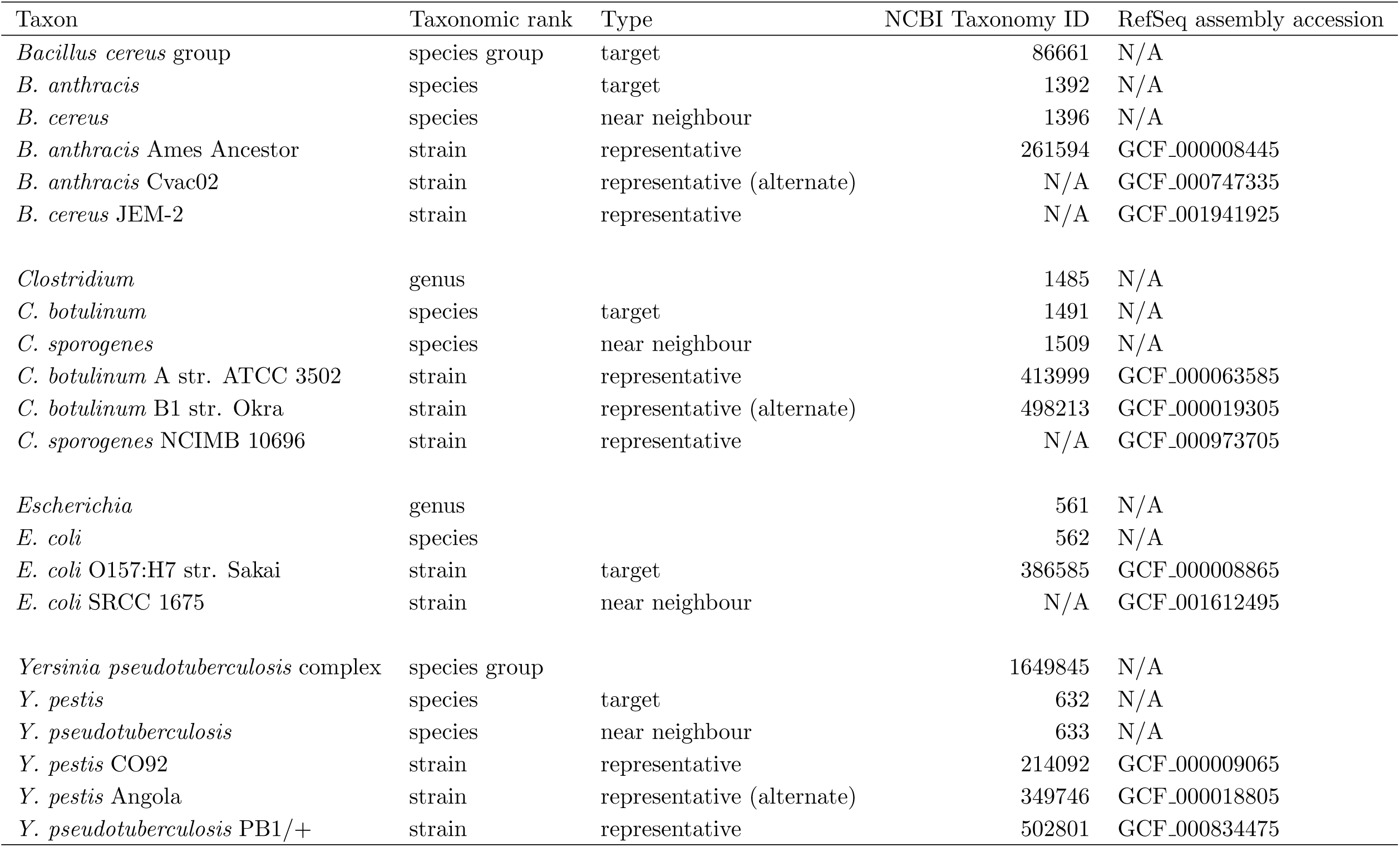
Taxonomic data and metadata for target taxa and near neighbour species or strains.

### Clade-level exclusion

In the final read simulation experiment, clade-level exclusion [9] was performed to assess classification performance in the situation where the taxon for which one has sequence data is not represented in the reference database. In these tests, we simulated 250-bp reads from the taxon hypothetically missing from reference database, excluded this taxon from the reference database when performing blast searches, and then obtained optimal filter parameter values for classification of the target taxon, which in this case was the taxon immediately above the excluded taxon in taxonomic rank.

## Results and discussion

### Evaluation of a “first-pass” taxonomic classifier

To demonstrate typical use of a first-pass taxonomic classification program, we analyzed all simulated reads from the *B. cereus* JEM-2 genome [34,35] with Kraken [36]. The majority of the reads (79%) were assigned to the *Bacillus cereus* group; of these, only 32% of the reads were assigned more specifically to *B. cereus*. Worryingly, however, a relatively small number of reads were assigned incorrectly to other *Bacillus cereus* group species, including *B. anthracis, B. cytotoxicus, B. mycoides, B. thuringiensis*, and *B. weihenstephanensis*. Had this benign strain of *B. cereus* (JEM-2) been the sole representative of the *Bacillus cereus* group in a metagenomic sample, an analyst using Kraken might have erroneously declared that a variety of *Bacillus cereus* group species were present in the sample, including pathogenic *B. anthracis*. As false-positive assignments are relatively commonplace with first-pass classification programs, we were motivated to develop a procedure to validate the assignments of interest made by such classifiers.

### Taxon selection

To demonstrate the blast-based validation procedure, we selected four target taxa, all of which are biological agents that could conceivably be of interest in a bioforensic setting. The first is *Bacillus anthracis*, the bacterium that causes anthrax. The second is *Clostridium botulinum*, a bacterium capable of producing the lethal botulinum neurotoxin. The third is a pathogenic strain of *Escherichia coli, E. coli* O157:H7 str. Sakai, a bacterium that has been associated with major outbreaks of foodborne illness. The fourth and final target taxon is *Yersinia pestis*, the bacterium that causes bubonic plague. Thus, three out of the four target taxa represent particular species to be identified (*B. anthracis, C. botulinum*, and *Y. pestis*), whereas one target taxon represents a particular strain to be identified (*E. coli* O157:H7 str. Sakai). Species-level evaluations were performed using the representative strains indicated in Table 1. In two of the three evaluations, the genome chosen was a reference genome for the species (C. *botulinum* A str. ATCC 3502 and *Y. pestis* CO92). For the *B. anthracis* evaluation, the genome of the Ames Ancestor strain was used to ensure that the pXO plasmids were included, as presence of the pXO plasmids is normally required for *B. anthracis* to be fully virulent [24, 25, 29]. To evaluate the implications of representative genome choice, an alternate representative genome was selected for each species. Additional information about the target taxa and near neighbours is provided in Table 1.

### Simulated read experiments

A total of four simulated read experiments were performed to determine optimal blast filter parameter values for the identification of various target taxa. A comparison of sensitivity across experiments on a per-taxon basis is available in Supplementary Figs. S1-S4, online.

### Experiment 1: 250-bp simulated reads

The first experiment simulated 250-bp reads from the target and near neighbour genomes; the results are shown in Table 2.

**Table 2.** Experiment 1: simulated 250-bp reads from four target taxa. Optimal parameter values for filtering blast results were chosen to maximize precision (first) and sensitivity (second) using two different approaches to compute false positives.

| Target taxon | Taxonomic rank | Number of reads | Approach used to compute $FP$ | Maximum E-value | Bit score difference | Validated reads | Sensitivity | Precision |
| --- | --- | --- | --- | --- | --- | --- | --- | --- |
| *B. anthracis* | species | 220,140 | near neighbour | $10^{-64}$ | 8 | 19,491 | 0.088539 | 1.0 |
| | | | false negatives | $10^{-64}$ | 8 | 19,491 | 0.088539 | 0.999995<sup>1</sup> |
| | | | false negatives | $10^{-64}$ | 128 | 1,751 | 0.007954 | 1.0 |
| *C. botulinum* | species | 156,120 | near neighbour | $10^{-64}$ | 8 | 153,786 | 0.985050 | 1.0 |
| | | | false negatives | $10^{-64}$ | 8 | 153,786 | 0.985050 | 1.0 |
| *E. coli* O157:H7 str. Sakai | strain | 223,760 | near neighbour | $10^{-64}$ | 1 | 838 | 0.003745 | 1.0 |
| | | | false negatives | $10^{-64}$ | 8 | 184 | 0.000822 | 1.0 |
| *Y. pestis* | species | 193,170 | near neighbour | $10^{-64}$ | 8 | 20,398 | 0.105596 | 1.0 |
| | | | false negatives | $10^{-64}$ | 8 | 20,398 | 0.105596 | 1.0 |

<sup>1</sup> In the case of *B. anthracis*, we observe that sensitivity increased from $\approx 0.8\%$ to $\approx 8.9\%$ when allowing exactly one $FN$ assignment (precision $\approx 99.9995\%$ ).

We observe that when requiring perfect precision, sensitivity was highest for identification of *C. botulinum* (≈99%), followed by much lower sensitivity for *B. anthracis* and *Y. pestis*. These results are understandable, as it is well established that the species that comprise *B. cereus sensu lato* have very similar genomic content [6], and that *Y. pestis* and *Y. pseudotuberculosis* are also very closely related [1]. Sensitivity was lowest for identification of *E. coli* O157:H7 str. Sakai (≈0.4% for near neighbour and ≈0.08% for false negatives). Again, this result is consistent with the expectation that strain-level identification would be substantially more challenging than species-level identification, as the two *E. coli* strains in this case are ≈99.97% identical. Because the reads in this experiment were simulated from genomes that were present in the reference database, almost all read alignments had equally good scores, so the absolute E-value filter had little or no effect until it was set so stringently that it eliminated all *TP* (*E* = 10^−128^). In the case of *B. anthracis*, we observe that sensitivity increased from ≈0.8% to ≈8.9% when allowing exactly one FN assignment (precision ≈99.9995%; Table 2). This suggests that if one is willing to relax the perfect precision requirement very slightly, it may be possible to make significant gains in sensitivity. Finally, it is interesting to note that in most cases, *b* =8 maximized sensitivity while achieving perfect precision. This likely represented a “sweet spot” (at least as compared to *b* = 4 or *b* = 16) for the level of taxonomic specificity represented by the selected target taxa.

### Experiment 2: 250-bp simulated reads, alternate representative genome

Choosing a particular genome to represent a strain, species, or higher-level taxon could in principle have implications for the filter parameter values recommended by the optimisation procedure. While hopefully the taxonomy is structured such that members of a particular clade are more similar to each other than to members of other clades, taxonomies are well known to be imperfect in this regard. To test the implications of representative genome choice, we repeated the species-level evaluations from Experiment 1, except that we used an alternate representative genome for the target taxon, the database genome that was most distantly related to the original representative genome. The results are shown in Table 3.

**Table 3.** Experiment 2: simulated 250-bp reads from three target taxa using alternate representative genomes. Optimal parameter values for filtering blast results were chosen to maximize precision (first) and sensitivity (second) using two different approaches to compute false positives.

| Target taxon | Taxonomic rank | Number of reads | Approach used to compute $FP$ | Maximum E-value | Bit score difference | Validated reads | Sensitivity | Precision |
| --- | --- | --- | --- | --- | --- | --- | --- | --- |
| *B. anthracis* | species | 209,080 | near neighbour | $10^{-64}$ | 8 | 20,114 | 0.096202 | 1.0 |
| | | | false negatives | $10^{-64}$ | 8 | 20,114 | 0.096202 | 1.0 |
| *C. botulinum* | species | 164,270 | near neighbour | $10^{-64}$ | 8 | 162,069 | 0.986601 | 1.0 |
| | | | false negatives | $10^{-64}$ | 4 | 162,594 | 0.989797 | 1.0 |
| *Y. pestis* | species | 187,470 | near neighbour | $10^{-64}$ | 8 | 22,576 | 0.120425 | 1.0 |
| | | | false negatives | $10^{-64}$ | 8 | 22,576 | 0.120425 | 1.0 |

In general, the optimal parameter values recommended by this experiment and the resulting values of sensitivity and precision were highly concordant with the results of Experiment 1 (Tables 2 and 3). The optimal parameter values recommended for classification of *B. anthracis* when using the Cvac02 strain were identical to those recommended when using the Ames Ancestor strain (*E* = 10^−64^ and *b* =8), with the exception that it was possible to achieve perfect precision using the false negatives approach when *b* = 8. Likewise, when using *C. botulinum* B1 str. Okra, the false negatives approach recommended *b* = 4 rather than *b* = 8. These results suggest that the filter parameter values recommended by the false negatives approach are potentially more dependent on representative genome choice than those recommended by the near neighbour approach. The calculation of *FP* in the false negatives approach is based solely on the classification of reads simulated from the chosen representative genome, whereas in the near neighbour approach, *FP* can result from the assignment of a near neighbour read to any genome associated with the target taxon (any strain of *B. anthracis*, for example). Thus, it might behoove a user of our method to sample the diversity in their clade of interest by running the optimisation procedure for multiple representatives and using the globally most conservative recommended parameter values for classification (if maximizing precision is the goal). Alternatively, one might devise a method for more exhaustive sampling of the diversity that might exist among target taxon genomes.

### Experiment 3: 150-bp simulated reads

Experiment 3 was identical to Experiment 1, except that a simulated read length of 150 bp was used, thus making the classification task more difficult. The results are shown in Table 4.

**Table 4.** Experiment 3: simulated 150-bp reads from four target taxa. Optimal parameter values for filtering blast results were chosen to maximize precision (first) and sensitivity (second) using two different approaches to compute false positives.

| Target taxon | Taxonomic rank | Number of reads | Approach used to compute $FP$ | Maximum E-value | Bit score difference | Validated reads | Sensitivity | Precision |
| --- | --- | --- | --- | --- | --- | --- | --- | --- |
| *B. anthracis* | species | 366,920 | near neighbour | $10^{-32}$ | 8 | 16,904 | 0.046070 | 1.0 |
| | | | false negatives | $10^{-32}$ | 8 | 16,904 | 0.046070 | 1.0 |
| *C. botulinum* | species | 260,200 | near neighbour | $10^{-32}$ | 8 | 250,915 | 0.964316 | 1.0 |
| | | | false negatives | $10^{-32}$ | 8 | 250,915 | 0.964316 | 1.0 |
| *E. coli* O157:H7 str. Sakai | strain | 372,960 | near neighbour | $10^{-64}$ | 4 | 709 | 0.001901 | 1.0 |
| | | | false negatives | $10^{-64}$ | 8 | 180 | 0.000483 | 1.0 |
| *Y. pestis* | species | 321,970 | near neighbour | $10^{-32}$ | 8 | 19,965 | 0.062009 | 1.0 |
| | | | false negatives | $10^{-32}$ | 8 | 19,965 | 0.062009 | 1.0 |

With optimal filter parameter values, we observe that sensitivity in detecting each target taxon decreased relative to the 250-bp experiment — e.g., in the case of *C. botulinum*, sensitivity decreased from ≈99% to ≈96% (Tables 2 and 4). Also, optimal values for the E-value and bit score difference filters varied somewhat relative to the 250-bp experiment, although it was always the case that *E* ≤ 10^−64^ and *b* ≤ 8.

### Experiment 4: clade-level exclusion, 250-bp simulated reads

In a final simulated read experiment, clade-level exclusion of either species (*B. anthracis*) or strains (*C. botulinum* A str. ATCC 3502 and *Y. pestis* CO92) was performed to assess classification performance when the taxon for which one has sequence data is not represented in the reference database, a situation commonly encountered in practice. Only the false negatives method of computing *FP* was used; the results are shown in Table 5.

**Table 5.** Experiment 4: simulated 250-bp reads from three taxa that were summarily excluded from the reference database. Optimal parameter values for filtering blast results were chosen to maximize precision (first) and sensitivity (second) using the “false negatives” approach to compute false positives.

| Target taxon | Taxonomic rank | Excluded taxon | Number of reads | Maximum E-value | Bit score difference | Validated reads | Sensitivity | Precision |
| --- | --- | --- | --- | --- | --- | --- | --- | --- |
| *B. cereus* group | species group | *B. anthracis* | 220,140 | $10^{-64}$ | 64 | 36,003 | 0.163546 | 0.939339 |
| *C. botulinum* | species | *C. botulinum*<br>A str. ATCC 3502 | 156,120 | $10^{-64}$ | 32 | 141,277 | 0.904926 | 0.999385 |
| *Y. pestis* | species | *Y. pestis* CO92 | 193,170 | $10^{-64}$ | 8 | 20,272 | 0.104944 | 1.0 |

In this experiment, we observe that it was not always possible to achieve perfect precision — maximum precision for identification of the *B. cereus* group when excluding *B. anthracis* was ≈93.9%, and maximum precision for identification of *C. botulinum* when excluding *C. botulinum* A str. ATCC 3502 was ≈99.9%. We note that sensitivity for identification of *C. botulinum* decreased from ≈98.5% in Experiment 1 (Table 2) to ≈90.5% in the clade-level exclusion experiment (Table 5). By contrast, sensitivity for identification of *Y. pestis* hardly decreased at all (≈10.6% vs. ≈10.5%).

### Calculating precision: “near neighbour” versus “false negatives”

Our results show that it was possible to achieve perfect precision in 3/4 simulated read experiments when using either the near neighbour or the false negatives approach (the exception being the clade-level exclusion experiment). In 7/11 cases, the filter parameter values recommended by the two approaches were identical; in the cases where they differed, the false negatives approach uniformly recommended more stringent filter parameter values than the near neighbour approach, resulting in reduced sensitivity (Tables 2, 3, and 4). As mentioned previously, deciding which of the two approaches to calculating precision is superior would require a comparison to a provably optimal approach, which we currently deem computationally intractable. Each heuristic makes assumptions that may not always hold: the near neighbour approach assumes that a single genome that is closely related to the target taxon is sufficient to serve as a proxy for all other non-target taxa in the database, and the false negatives approach assumes that the sequencing error process is symmetric. When seeking to avoid an erroneous claim that a particular biological agent is present in a sample, one may wish to use the more conservative set of parameter values recommended by the two approaches.

### Practical application of the BLAST-based validation procedure

To demonstrate the practical application of the blast-based validation procedure, we downloaded a subset of metagenomic data collected from the New York City subway system (ncbi sra id SRR1748708), which the original study indicated might contain some reads from *B. anthracis* [2]. Indeed, analysis of this data with Kraken, our first-pass classifier, assigned 676 reads to *B. anthracis* (≈0.04% of reads). However, blast-validation of these 676 reads using the most conservative parameters recommended by our study (E = 10^−64^ and b = 128; Table 2) resulted in zero reads assigned to *B. anthracis*. Even after significantly relaxing the minimum required bit score difference (setting b =8), which was shown in Experiment 1 to significantly increase sensitivity (Table 2), still zero reads were assigned to *B. anthracis*. Thus, we would conclude that the 676 reads that Kraken assigned to *B. anthracis* were in fact false-positive assignments, which agrees with other follow-up studies that have been performed on the New York City subway data [13].

## Conclusions

We have shown how blast, a very widely used tool for sequence similarity searches, can be used to perform taxonomic assignment with maximal precision by using blast result filters fine-tuned via read simulation experiments in conjunction with an lca algorithm. We demonstrated the parameter optimisation process for four different pathogenic organisms, and showed that optimal parameter values and resulting values of sensitivity and precision varied significantly depending on the selected taxon, taxonomic rank, read length, and representation of the sequenced taxon in the reference database. Furthermore, the addition or removal of a single sequence from the reference database could change the recommended optimal parameter values, so the optimisation process should be re-run every time the database is updated.

Once optimal blast filter parameter values for a particular taxon have been determined, they can be subsequently used to perform validation of sequence assignments to that taxon. Given the massive size of many metagenomic data sets, however, we envision most users employing a “two-step” approach that involves first producing candidate target taxon sequence assignments using a relatively fast classification program — one that is not necessarily optimised for precision — and then confirming the veracity of those sequence assignments using the blast-based validation procedure.

One would be hard-pressed to define a “typical” metagenomic experiment, and the probability that a particular genome that is physically present in a metagenomic sample at some abundance is ultimately represented in the sequencing library and sequenced to a particular degree of coverage is a function of many factors that are outside the scope of this study. The methods we present here are concerned with read-by-read taxonomic assignment (each read interrogated independently of all other reads), and the selection of optimal blast result filters for this assignment process — in our case, we define “optimal” to mean correct assignment of the greatest possible number of reads without any incorrect assignments. In a real-world detection scenario, an additional question will often be asked: how many reads should be assigned to a particular target taxon before one deems it “present” in the sample? In principle, if one assumes that the reads in question originate from a genome that is present in the reference database, and that there was no error associated with the read simulation process or choice of optimal filter parameter values, then the answer is simply “one read”. In practice, however, if only one read out of millions or billions is assigned to a particular taxon, it is only natural that one may hesitate to claim that a potential pathogen or other biological agent is present in a sample on the basis of such scanty evidence. Unfortunately, meaningful additional guidance on this point would require a comprehensive accounting of all possible sources of error associated with the analysis of a metagenomic sample.

Potential users of the software will find scripts for parsing blast results, performing parameter sweeps, and assigning final taxon labels to sequences at https://github.com/bioforensics/blast-validate.

## Acknowledgements

We thank Todd Treangen and Nicholas Bergman for helpful discussions about the project, and Travis Wyman, Timothy Stockwell, and Lisa Rowe for providing feedback on drafts of the manuscript.

## Additional Information and Declarations

### Author contributions

ALB conceived the experiments and all authors participated in the intellectual development of the project. BDO, DDS, and ALB wrote the software. All authors took part in conducting the experiments and analysing the results. ALB drafted the manuscript and all authors reviewed and revised the text.

### Software and data availability

All genome data used in this study is available from the ncbi RefSeq database [26] (assembly accessions provided in Table 1). The software implementing the methods described in this study, as well as simulated reads, analysis results, and other files germane to this study are available online at the following url: https://github.com/bioforensics/blast-validate

